# Rutin, a Natural Inhibitor of IGPD Protein, Inhibits the Biofilm Formation in *Staphylococcus xylosus* ATCC700404

**DOI:** 10.1101/802447

**Authors:** God’spower Bello-Onaghise, Xing Xiaoxu, Zhou Yonghui, Qu Qianwei, Cui Wenqiang, Tang Yang, Chen Xingru, Wang Jinpeng, Yan-Hua Li

## Abstract

The biofilm of bacteria plays an important role in antibiotic resistance and chronic infection. Thus, in order to solve the problem of resistant bacteria, it is very important to find new drugs that can inhibit the formation of biofilms. In recent years, researchers have shifted their attention to natural products. As a flavonoid, rutin has been reported to have a variety of biological activities, interestingly, in this study, the inhibitory effect of rutin on the biofilm of *Staphylococcus xylosus* was investigated. We confirmed that rutin could effectively inhibit the biofilm formation of *S. xylosus*, then, for the sake of discussion on how it interferes with the biofilm formation, the interaction between rutin and imidazolyl phosphate dehydratase (IGPD) which has been identified as the key enzyme that plays a vital role in the process of biofilm formation was analyzed by molecular docking, the results showed that rutin had a strong affinity with IGPD, it occupied the hydrophobic cavity of the active center forming four hydrogen bonds and many other interactions. In addition, we proved that rutin was able to combine with IGPD using SPR technique. Therefore, we determined the enzyme activity and histidine content of IGPD, the result indicated that rutin could simultaneously inhibit the activity of IGPD and abrogate the synthesis of histidine. Interestingly, the *hisB* gene encoding for IGPD and IGPD in *S. xylosus* were also significantly inhibited when the bacterial culture was treated with rutin. Taken together, the results have provided evidence that rutin is a natural drug that has the ability to interfere with the formation of biofilm in *S. xylosus*. It is therefore a potential enzyme inhibitor of IGPD.

**Author’s Summary:** *Staphylococcus xylosus* has been isolated from a variety of infections, and the biofilm formed by *S. xylosus* can help the bacteria evade the immune system of the host and cause chronic infections. Here, we dealt with this menace by establishing a highly effective drug with the ability to interfere with the process involved in the formation of biofilm in *S. xylosus*. IGPD has been reported to be directly involved in the formation of biofilm in *Staphylococcus xylosus* and it is known to be present in a variety of microorganisms. Based on this study, we developed a drug therapy targeting IGPD and at the same time interfere with the formation of biofilm in *S. xylosus*

## INTRODUCTION

*Staphylococcus xylosus* is a coagulase-negative, gram positive coccoid organism that was first identified in 1975 [1]. *S. xylosus* are common bacteria in the environment and have been linked to opportunistic infections in both humans and animals. Although there are few reports on the pathogenicity of *S. xylosus*, there have been some cases of *Staphylococcus xylosus* infection reported in recent years [2], including erythema nodosum [3], pyelonephritis[4], mastitis [5] and corneal infections [6]. In addition, *S. xylosus* was also isolated from the posterior coronal sulcus of male circumcision cases [7]. This has drawn the attention of researchers and clinicians to the potential public health threat of S*. xylosus*. What is more serious is that, S*. xylosus* has a strong ability to form biofilm [8, 9]. Biofilms have been defined as multiplex networks consisting of a consortium of bacteria surrounded by a self-produced three-dimensional organic extracellular matrix [10–15] and bio-minerals [16, 17]. The formation comprises a number of well-organized steps including attachment to a surface, formation of microcolonies, formation of young biofilm, development of differentiated structures in which individual bacteria as well as the entire community are surrounded by exopolysaccharides[14], and subsequent dispersal of mature biofilm[18].The dispersal of mature biofilms marks the completion of a full circle[19]. Biofilms provide an effective protective cover for the residing bacteria, making them evasive to their host immune systems and about 1000 times more resistant to environmental pressures and therapeutic interventions by antibiotics than their planktonic counterparts [14, 20, 21], leading to chronic infections [22]. About 80% of the world’s persistent and chronic infections have been reported to be mediated by bacteria that live within biofilms [23, 24]. Thus, the search for novel antibiotics with the ability to interfere with the process biofilm formation in bacteria is urgent and very critical. This will help in mitigating chronic and persistent infections mediated by biofilm forming bacteria such as *S. xylosus*.

Imidazoleglycerol phosphate dehydratase (IGPD) is the major enzyme in the pathway of histidine biosynthesis in bacteria and plants [25, 26]. As one of the pathways of nitrogen metabolism, L-histidine synthesis pathway is related to biofilm formation [27, 28]. IGPD catalyzes the sixth step of histidine biosynthesis which involves a dehydration reaction to produce imidazoleacetol phosphate (IAP) from imidazoleglycerol phosphate (IGP) and a concomitant water molecule [29–32], it is often used as a target for herbicides because of its unique biological characteristics [26, 33]. The presence of two molecules of metal ion Mn^2+^ at the active center of IGPD plays an important role in catalyzing substrate reaction [32]. These indicate the direction of IGPD inhibitors. In a previous study, we demonstrated that IGPD plays an important regulatory role in the biofilm formation of *S. xylosus* [34]. Therefore, IGPD was considered to be a potential target in solving the menace of *S. xylosus.* The spatial structure of IGPD was predicted by homologous module, and it was used as the basis for screening the enzyme inhibitor [35].

Rutin, also known as vitamin P is a major flavonoid. It has been reported to possess antiplatelet, antiviral and antihypertensive properties. It also enhances the functions of blood capillaries because of its high cardioprotective and antioxidant activity [36]. In addition, the antibiofilm activity of rutin has also been reported. For example, data obtained from betabolomics-based screening revealed that burdock leaf significantly inhibited the formation of biofilm by *Pseudomonas aeruginosa* due to its high rutin content [37]. This was further corroborated by findings from other authors. It was observed that rutin obtained from aqueous extract of *syringa oblata lindl* inhibited the formation of biofilm by *streptococcus suis* [38, 39], *Escherichia coli* and *Staphylococcus aureus* [40]. However, the effect of rutin on the biofilm of *S. xylosus* has not been investigated.

Furthermore, some recent reports have also described rutin as a protease inhibitor [41]. Ragunathan and Ravi have used molecular docking techniques to suggest that rutin may be a potential inhibitor of protein [2]. So far, there is no report on the inhibitory effect of rutin on the proteins that promote the formation of biofilms. In this study, we evaluated the effect of rutin on *S. xylosus* biofilm. We predicted the potential binding mode of rutin and IGPD by molecular docking and molecular dynamics. Then, we used surface plasma resonance, enzyme activity analysis and amino acid site directed mutagenesis to verify the interaction between rutin and IGPD.

## RESULTS

### Determination of the ability of rutin to inhibit the biofilm formation of *Staphylococcus xylosus* (ATCC700404)

In order to determine whether rutin could interfere the formation of biofilm of *S. xylosus* we selected different concentrations of rutin which were dissolved in the 10 % (v/v) methyl alcohol to conduct the assay. Although, there was no statistical significant inhibitory effect on the growth of *S. xylosus* in the experiment (as shown in Fig 1 B), 0.8mg/ml of rutin had no effect on the growth rate of *S. xylosus* at 0 - 12 hours at 37°C), however, a closer look at the result reveals an interesting trend of decrease in growth. In comparing the behavior of strains without rutin (control) with those treated with the different concentrations (0.2mg/mL, 0.4mg/mL, 0.8mg/mL) of rutin, there was a dose-dependent growth reduction effect. The strains treated with 0.8mg/ml rutin had a significantly lower (p < 0.05) growth rate than others (see Fig 1A). On the other hand, the nature and structure of the biofilm was not significantly affected following growth in the presence of 0.2mg/ml, 0.4mg/ml rutin (p > 0.05), however, 0.8 mg/ml of rutin had a significant inhibitory effect on the formation of biofilm by *S. xylosus* after incubation for 3 hours, 6 hours 12 hours and 24 hours (p<0.05), respectively.

**Fig 1:**
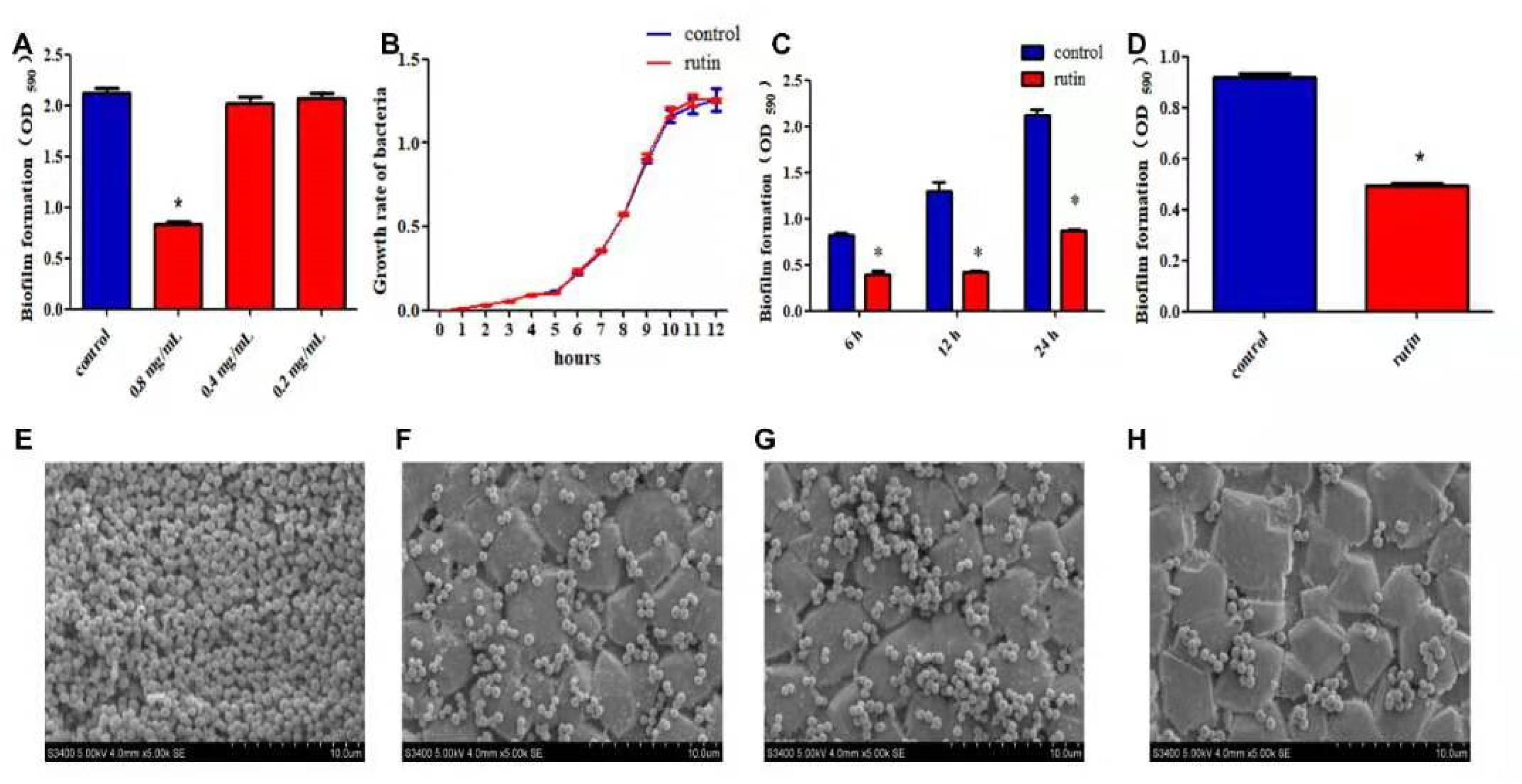
Inhibitory effect of Rutin on the biofilm formation of *S. xylosus ATCC700404*. A: Inhibitory effect of Rutin with different concentrations on the biofilm formation of *S. xylosus ATCC700404*, 0.8mg/ml rutin significantly inhibited the formation of biofilm (p<0.05), but the other two concentrations did not. (p>0.05). B: The effect of rutin on the growth rate of *S. xylosus.* C: The effect of 0.8 mg/ml rutin on the biofilm of *S. xylosus* at different growth stages. When the bacteria were incubated for 6, 12, and 24 hours, respectively, rutin significantly inhibited the formation of the biofilm (p<0.05). D: Rutin inhibited the formation of the biofilm of *ΔhisB S. xylosus* at 24 h. Scanning electron micrographs of *S. xylosus* ATCC700404 biofilm following growth in TSB supplemented with 0.8mg/ml of rutin (F), or control (E). Controls refer to the cultures without rutin. Scanning electron micrographs of *ΔhisB S. xylosus* biofilm following growth in TSB supplemented with 0.8mg/ml of rutin (H), or control (G). Controls refer to the cultures without.

In addition, we also measured the effect of rutin on the biofilm of *ΔhisB S*. *xylosus*. The experimental results showed that rutin also had an intervention effect on the biofilm of *ΔhisB S*. *xylosus*, as shown in Fig 1D. In order to further verify the intervention of rutin on the biofilm of *S*. *xylosus* and *ΔhisB S*. *xylosus*, we observed the effect of rutin on the biofilm morphology by scanning electron microscopy. As shown in Fig 1(E - H, a thick structure of biofilm made up of aggregates and micro colonies of *S*. *xylosus* almost completely covered the surface of the rough glass slide (control; see Fig 1E), however, when the culture medium was supplemented with rutin (0.8 mg/ml), only a small amount of *S*. *xylosus* was attached on the glass (Fig 1F). Interestingly, only a small number of *ΔhisB S*. *xylosus* were deposited on the glass plate (Fig 1G), although it was observed that the number of *ΔhisB S*. *xylosus* decreased significantly after adding 0.8 mg/ml rutin (Fig 1H).

### Interaction between Rutin and IGPD

First, our computational approach was initiated by docking of the 3D structure of rutin into the binding pocket of the active structure of IGPD. The results of the docking study revealed that rutin binds to the IGPD in a highly favorable manner in all of the predicted interaction patterns (Fig 2A). As many as nine amino acid residues and rutin formed ligand receptor interactions. Among them, Glu11, lle27, His36, Glu58 and rutin formed a strong hydrogen bond respectively (see Fig 2B). Interestingly, the bonds formed with IGPD were evenly distributed around rutin so that it could be firmly fixed in the active region of IGPD. On this basis, we carried out the simulation of molecular dynamics. The RMSD value of the complex structure was also relatively stable in the calculation process, which proves that the molecular docking between rutin and IGPD was reasonable (Fig 2C). The results of ligand receptor interaction energy are shown in Fig 2D. In the process of simulating the whole molecular dynamics, the energy of the ligand receptor complex was continuously reduced. The energy value was -102.500 when the calculation was over. This showed that the interaction between rutin and IGPD is quite stable. The above simulation results show that IGPD is a potential target for rutin. To verify this claim, we carried out molecular interaction test and enzyme activity analysis test. SPR analysis shows that rutin can interact with IGPD. When rutin was dissociated, the interaction disappeared (Fig 2E). We also found that the interaction between rutin and IGPD increased as the concentration of rutin increases (Fig 2F). However, this only proves that rutin can bind to IGPD, and does not mean that rutin is an inhibitor of IGPD. The enzyme activity analysis experiment can provide a clue to understanding the inhibitory effect of rutin on IGPD. The bacteria strains were grown in media with and without (control) 0.8mg/ml rutin at the 6 hours, 12 hours or 24 hours. The reaction was significantly lower in treated medium compared with the control (p<0.05). The results showed that rutin inhibited the enzyme activity of IGPD (Fig 2 G).

**Fig 2:**
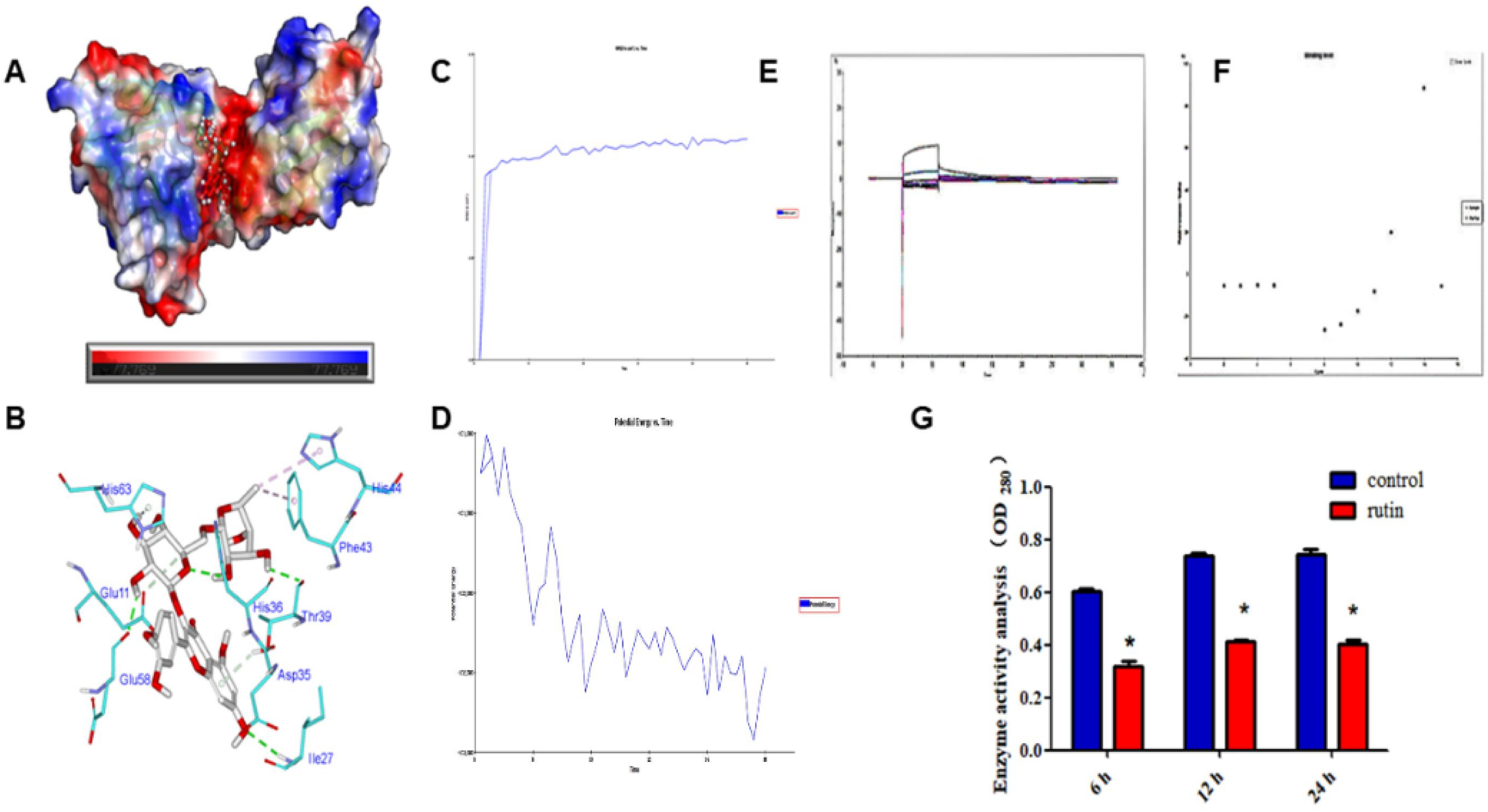
3D structure showing the docking of rutin with IGPD molecule. A: The 3D structure of rutin docking with IGPD molecule shows the distribution of energy and interaction. The red region represents the concentration of hydrogen bonds. B: Distribution of Interaction Forces between Rutin and Amino Acids around IGPD. C: Molecular dynamics of protein-ligand complexes formed by rutin and IGPD were calculated. RMSD value is in a relatively stable state from the beginning to the end of calculation. D: Interaction energy trends of protein ligand complexes in MD calculations. The interaction between rutin and IGPD was detected by SPR. When rutin was added to a chip with IGPD fixed on the surface, it generated signals representing the interaction (E). When the concentration of rutin was between ImM-10mM, the interaction intensity showed a positive correlation (F). G: Rutin was added before the growth. The bacteria grew at 6, 12 and 24 hours, and the protein activity of IGPD was extracted by ultrasonic sonicator. The activity of the enzyme was measured and compared with the enzyme activity of bacteria strains in culture without rutin.

### Identification of Rutin Binding Sites to IGPD

By analyzing the first four dominant conformations of rutin and IGPD, we found that both Ile27 and His36 formed hydrogen bonds with rutin. We suspected that these two amino acids play an important role in the binding of rutin and IGPD. Then we constructed three 3D models of mutant proteins and docked them with rutin separately. The results are shown in Fig 3 (A - C). It was observed that the interaction between rutin and the double mutant proteins is less than that between rutin and IGPD, and only one hydrogen bond is formed between rutin and double mutant protein (Fig 3C). This indicated that the change in the amino acid chain was not conducive to the binding of rutin to IGPD. Then we constructed and expressed the three mutant proteins, which were successfully expressed as shown in Fig 3E and we analyzed the effect of rutin on the activity of the three mutant proteases. As expected, the changes in the amino acids affected the activity of the enzyme (see Fig 3F), resulting in a decline in the activity of the enzyme, but the activity was not completely lost. Following this, the next step was to verify whether the mutant amino acid is a potential binding site for rutin which was added to a solution containing substrate and mutant protein. Interestingly, the protein activity of the two single point mutations (lle27, His36) were significantly reduced after the addition of 0.8mg/ml rutin (p<0.05). However, the enzyme activity of the protein obtained by the double point mutation (lle27 and His36) did not changed (p>0.05).

**Fig 3:**
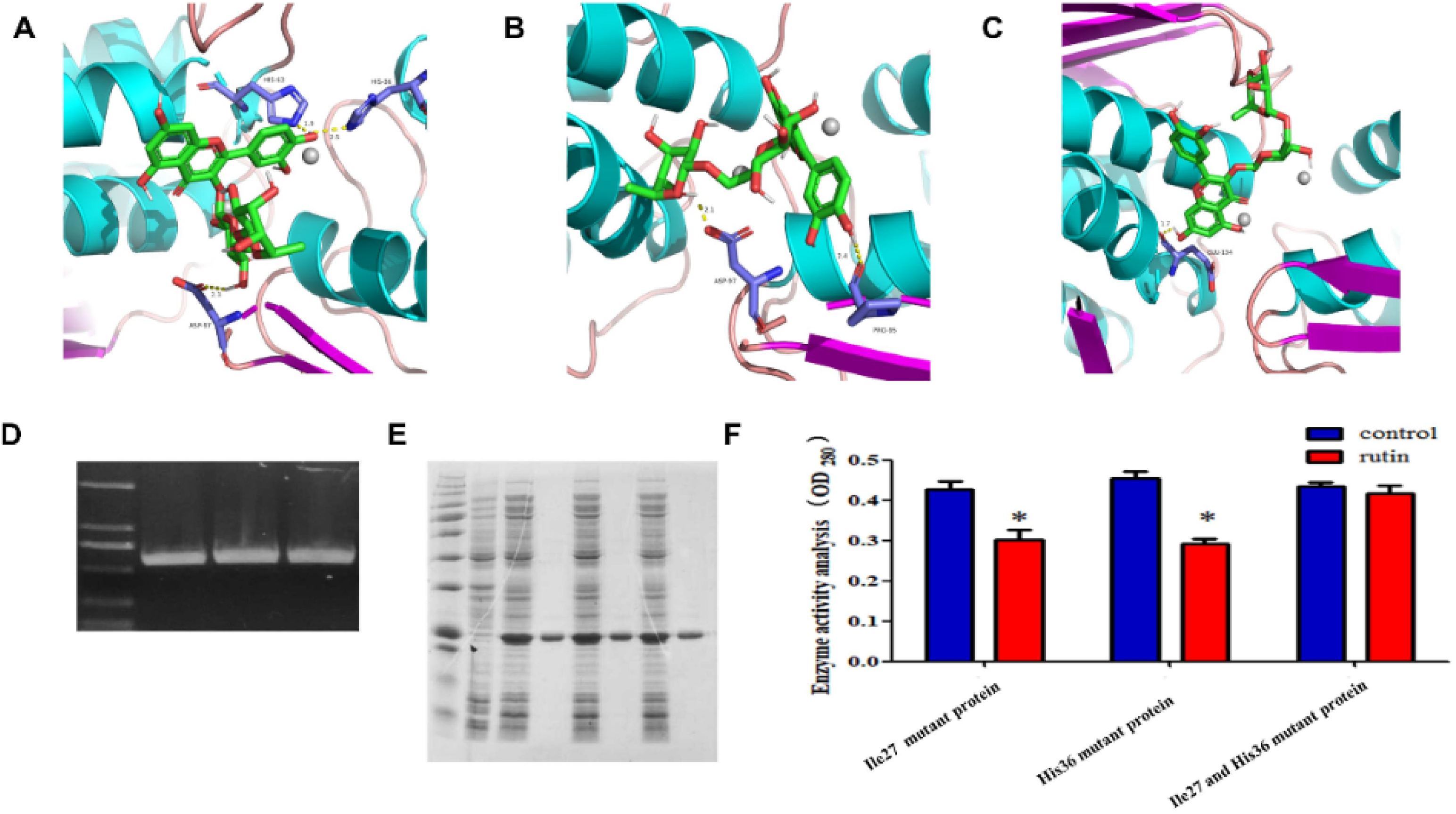
Molecular docking of the optimum conformation of rutin with the mutant proteins. A: Molecular docking of optimum conformation of rutin and Ile27 mutant protein. B: Molecular docking of optimum conformation of rutin and His36 mutant protein. C: Molecular docking of optimum conformation of rutin and double mutant protein (Ile27 and His36), D: The PCR results of three mutant genes were cloned successfully (From left to right are market, Ile27 single mutation, His36 single mutation, Ile27 and His36 double mutation). E: The results of the expression and purification of the three mutant proteins were shown through SDS-PEAG (From left to right: maker, Pre-expression bacterial supernatant, Pre-purified single mutant protein(lle27) supernatant, Purified single mutant protein (lle27)supernatant, Pre-purified single mutant protein(his36) supernatant, Purified single mutant protein (his36) supernatant, Pre-purified double mutant protein(lle27 and his36) supernatant and Purified double mutant protein (lle27 and his36) supernatant. F: Rutin was added to the solution containing the mutant protein, the control had no rutin, and the enzyme activity of each mutant protein was determined.

### Regulation of rutin on IGPD expression

First The effects of rutin on the *hisB* gene of *S. xylose* at 6h, 12h, and 24h, respectively, were measured by fluorescence quantitative PCR. The results show that 0.8mg/ml rutin could effectively reduce the amount of gene expressed at the various time points mentioned above (Fig 4A). Secondly, we measured the effect of rutin on the expression of IGPD protein. The results showed that rutin significantly reduce the expression of IGPD protein at 12 and 24 hours, but did not affect the expression of IGPD protein at 6 hours. See Fig 4B.

**Fig 4:**
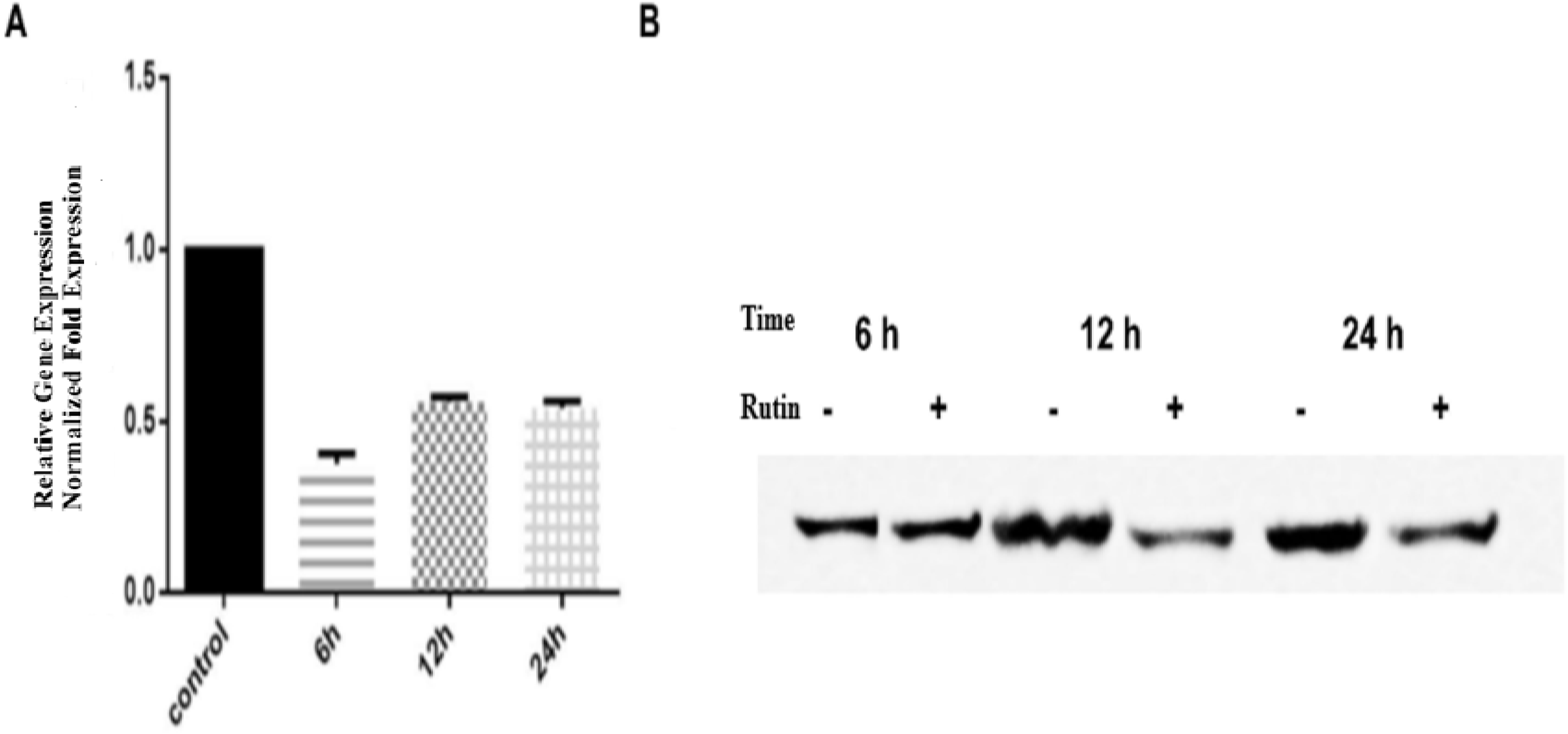
The effect of rutin on the expression of *hisB* gene in *S. xylosus*. A: The effect of rutin on the expression of *hisB* gene in *S. xylosus* at the 6, 12 and 24 h, respectively. B: The effect of rutin on the expression of IGPD protein in *S. xylosus* at 6, 12, and 24h, respectively.

### Determination of histidine content

In order to evaluate the nature and condition of IGPD, the histidine content of the bacterial strains was measured. This is because IGPD is an important protease in histidine synthesis pathway. As shown in Fig 5 (A), the histidine content of the bacteria growing in the medium treated with 0.8mg/ml rutin was significantly lower than that of the bacteria growing in the medium without the drug after 6 hours, 12 hours, and 24 hours(p<0.05), respectively.

**Fig 5:**
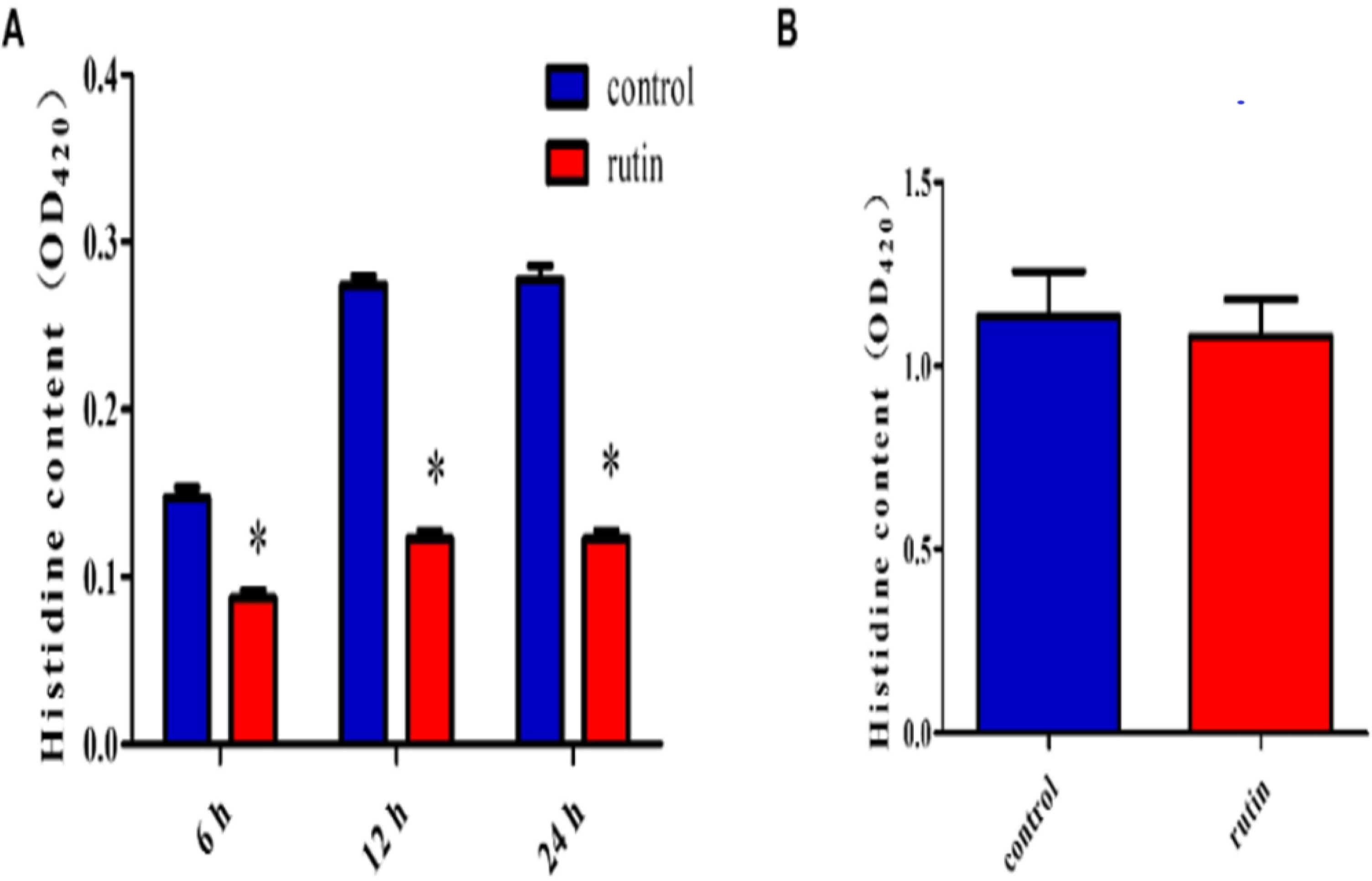
The determination of the effect of rutin on histidine content of *S. xylosus*. A: Rutin significantly reduced the histidine content of *S. xylosus* at 6, 12, and 24 hours, respectively of bacterial growth. B: The effect of rutin on the histidine content of the *ΔhisB S. xylosus* at the 24 h.

## DISCUSSION

Biofilms are consortia of bacteria that frequently dwell on medical devices, as well as environmental and biological surfaces [10–15]. Often, biofilms are comprised of diverse bacterial species that participate in synergistic interactions and contribute to recalcitrant infections [23, 24, 42]. In addition, bacteria living within a biofilm are typically more resistant to antimicrobials and have the ability to evade clearance by the host immune response [14, 20, 21, 43, 44]. *Staphylococcus xylosus* is widespread in the environment. Although no serious pathogenicity has been found in current studies, however, *S. xylosus* has been isolated in some infectious cases, such as erythema nodosum [3], pyelonephritis [4] and corneal infections [6]. Even if the role of *S. xylosus* in these infections is uncertain, the biofilm formed by *S. xylosus* makes these infections pathogenically more chronic and difficult to cure because under such biofilm protection, the bacteria have greater chance of prolong survival in the host, thus rendering the intervention of therapeutic drugs insignificant. Thus, there is an urgent need to find an alternative and more effective strategy to interfere with the process of biofilm formation in pathogenic bacteria such as *S. xylosus* so as to abrogate their pathogenesis in living hosts.

In this study, it was observed that 0.8 mg/ml rutin could effectively interfere with the formation of biofilm by *S. xylosus*. At the same time, we measured the effect of rutin on the formation of biofilm in *S. xylose* at 6, 12 and 24 hours, respectively. The results showed that rutin could significantly reduce the structure of biofilm of *S. xylosus* even as early as within 6 hours of intervention. This may be due to the fact that rutin can inhibit bacterial adhesion. According to earlier reports, rutin could interfere with the adhesion phase of *Streptococcus suis* and then abrogates the formation of its biofilm [45]. In addition, the bacterial quorum sensing system is also closely related to the early adhesion of bacteria. Besides, rutin could interrupt the production of AI-2 to inhibit the quorum sensing system of *Escherichia coli,* reduce bacterial adhesion, and then interfere with the formation of biofilm of *E. coli* [46]. It was reported that in combination with gentamicin sulfate, Rutin significantly weakened the adhesion of *Pseudomonas aeruginosa* and its ability to form biofilm [39]. Therefore, the strong inhibitory effect rutin had on the biofilm of *S. xylosus* at the different time points (6, 12, and 24 hours, respectively) in our study could be related to its ability to inhibit bacterial adhesion. Furthermore, the effect of rutin on the biofilm morphology of *S. xylosus* was observed by SEM. The results showed that rutin significantly destroyed the whole structure of the biofilm when the bacterial strains were incubated for 24 hours, a small number of the bacterial strains were seen growing independently on the slide. The results were consistent with those of crystal violet staining method, which further proved that rutin could destroy the biofilm structure of *S. xylosus*.

Molecular docking has become a new way to screen target protein inhibitors. Through molecular docking and virtual screening, Gruneberg [47] successfully identified several carbonic anhydrase inhibitors. Rastelli and co-workers [48] screened new aldose reductase inhibitors from the original database through molecular docking. Previous studies have demonstrated that IGPD plays an important regulatory role in the formation of biofilm in *S. xylosus* [34]. At the same time, we obtained the 3D structure of IGPD of *S. xylosus* by homologous modeling [35]. We attempted to predict whether rutin can bind to the binding sites of IGPD by molecular docking, so as to elucidate how rutin interfered with the biofilm formation process in *S. xylosus.* From the result of the molecular docking of rutin with IGPD, we observed that the scores of the first four binding conformations were above 100, and the Gibbs free energy was less than zero, which indicated that the binding of rutin with IGPD was spontaneous. In the optimal binding conformation of rutin to IGPD, rutin interacts with up to nine amino acids, including four hydrogen bonds and four Pi (∏) bonds. This strong interaction enables rutin to be effectively immobilized in the active region, suggesting that rutin is a potential inhibitor of IGPD [35]. In the subsequent calculations, we used the relevant program of molecular dynamics, which can more accurately reflect the state of receptor-ligand complexes than molecular docking. It has proven to be able to study a wide range of phenomena related to simple and complex molecules [49]. In this study, the protein ligand complexes obtained from the molecular docking procedure were calculated by molecular dynamics. The results revealed that the interaction energy between rutin and IGPD decreases continuously over time. In addition, RMSD values showed that the atomic deviation was small and the whole system was more stable, indicating that rutin and IGPD can spontaneously combine. Firstly, Surface plasmon resonance (SPR), is a sensor-based technology that allows the interaction of biological molecules to be determined in real-time [50–53]. Our findings suggested that it made a binding signal on the surface of a protein bound chip when 1 nm rutin was injected. In addition, this combination is dependent on concentration, when the concentration of rutin was between 1nm and 10nm, the interaction intensity showed a positive correlation (Fig 2F). This indicates that rutin can interact with IGPD [54]. Secondly, we determined the inhibitory effect of rutin on IGPD by enzyme activity assay. The results showed that the presence of rutin had a significant effect on the catalytic rate of the enzyme, however, we did not determine the half maximal inhibitory concentration (IC_50_) of rutin, because the main focus of the study was only on the concentration with the most effective and significant biofilm inhibition effect.

Looking at the interaction between rutin and the amino acid residues in the active chamber, it can be observed that the hydrogen bond formed between rutin and His36 was short, strong and very firm, followed by the hydrogen bond formed between rutin and Ile27. It was speculated that these two amino acids were the key amino acids that mediated the interaction of rutin with IGPD. Three 3D structures of mutant proteins, including Ile27 single mutant protein, His36 single mutant protein and Ile27 and His36 double mutant protein, were obtained. The mutant amino acids were replaced by alanine and docked with rutin in turn. The results showed that rutin still occupied the active cavity of the mutant protein (but with the double mutant protein). From the result of protein docking, it was observed that rutin only formed a hydrogen bond, and the score showed that the result of the docking was only 66.9. To verify the result of molecular docking, two-unit point mutation proteins and a two-site mutation protein were obtained by amino acid site-directed mutagenesis. Alanine was used to replace the mutant amino acid so as to ensure that the structure and function of the protein is not lost or significantly changed [55]. Next, the effect of rutin on the activity of mutant proteases was detected by enzyme activity analysis. The results showed that rutin could not inhibit the activity of IGPD when the two loci changed simultaneously. Amino acid mutation can change the structure of the active cavity, which may cause the radius of the active cavity to increase [55]. This made rutin to swing more freely in the active cavity, the instability of rutin in the active cavity is not be conducive for the interaction between rutin and the surrounding amino acids, and cannot effectively occupy the active sites, thus the inhibition of IGPD impossible. In addition, the mutation of two amino acids is essential for the formation of interaction force with rutin. When the two amino acids change, the distribution of the electron cloud of amino acid residues in the active cavity may change to some extent, which may be unfavorable for rutin binding. Interestingly, the two amino acids that could interact strongly with rutin were replaced at the same time. This may be the main reason why rutin could not bind to IGPD.

IGPD plays an important role in the formation of biofilm of S. xylosus [34]. Therefore, it was speculated that rutin may regulate the formation of biofilm in *S. xylosus* by regulating the transcription and translation levels of *hisB* gene leading to changes in the L-histidine content and enzyme activity. In order to verify this, we measured the regulation of rutin on *hisB* gene at different time points by RT-PCR. The results showed that rutin significantly reduced the expression of *hisB* gene. In order to further confirm this effect, we detected the effect of rutin on the expression of IGPD protein at different time points by Western blot. The result proved that rutin significantly reduced the protein expression of IGPD. However, gene and protein expression levels changed at different time points, which may be related to the differences in the gene levels caused by the different stages of bacterial growth [56]. In addition, we observed that there was significant difference between the gene expression and protein level at 6 hours. Rutin could reduce gene expression, but had no effect on the protein level at this stage. This may be due to a variety of factors, including low level protein expression, sample processing, and the actual relationship between transcript and protein abundance [57–63].

IGPD is one of the most important metalloenzymes in L-histidine biosynthesis [29–32]. In our study, 0.8mg/ml rutin significantly inhibited the activity of IGPD, which may affect the biosynthesis of L-histidine. So, in order to verify this claim, we used ultraviolet spectrophotometer to determine the effect of rutin (0.8mg/ml) on the histidine content of *S. xylosus* at 6, 12 and 24 hours, respectively. The effect of rutin was consistent with the expected results. Rutin significantly reduced the content of L-histidine at different time points. However, we also postulated that the content of L-histidine may also be affected by the metabolic pathways. Again, to further substantiate this, we examined the effect of rutin (0.8mg/ml) on the content of L-histidine in the *ΔhisB S. xylosus*. The results showed that there was no significant difference between the two groups. This indicated that the effect of rutin on the L-histidine content of *S. xylosus* was only caused by its interference with the activity of IGPD and thereby inhibiting its biosynthesis.

As we have previously analyzed, rutin inhibits the activity of IGPD by binding to the active site of IGPD, which leads to a decrease in L-histidine biosynthesis, thereby interfering with the formation of biofilm by *S. xylosus*. In addition, L-histidine biosynthesis regulates the expression of *hisB* gene. When bacteria are deprived of L-histidine, their *hisB* gene level will rise to cope with this change [64]. However, the results of the study showed that rutin inhibitd the activity of IGPD and the expression of *hisB* gene at the same time. As a result of this, the bacteria could not synthesize their L-histidine by themselves, thereby leading to a significant reduction in their histidine content.

In conclusion, rutin regulated the expression of IGPD, by binding to IGPD Ile27 and His36 sites, it affected the activity of enzymes and changed the content of L-histidine and in the process, interfered with the formation of biofilm in *S. xylosus*. This study is based on targeting IGPD as the major protein for drug intervention in the formation of biofilm by *S. xylosus*. From the foregoing therefore, the mechanism by which rutin interfered with the formation of biofilm in *S. xylosus* was well elucidated, and novel insights for controlling biofilms mediated persistent and chronic bacterial infections were provided. This will go a long way in enhancing our understanding of IGPD as a target for inhibitors.

## MATERIALS AND METHODS

### Bacterial strain and growth conditions

*Staphylococcus xylosus* ATCC700404 and *ΔhisB S. xylosus* [34] were the major strains used in this study. Bacteria were grown at 37°C in Tryptic Soy Broth (TSB). The cultures were used for the biofilm assays and other bacterial experiments.

### Determination of Minimum Inhibitory Concentration of Rutin itinerary

The effect of the minimum inhibitory concentration of rutin on the growth of *S. xylosus* was repeated three times, but no concentration of rutin was observed to significantly inhibit the growth of the bacteria. The growth curve of the bacteria was drawn to determine whether rutin had an effect on the growth of *S. xylosus*. Overnight culture of bacterial strains was grown at 37°C for 24 h, then TSB medium was used to dilute the bacterial cultures to 1×10^5^colony forming units/mL. This was followed by the addition of 0.8mg/ml rutin. The mixture was grown in an incubator for 12 h at 37°C and the absorbance was read with a UV spectrophotometer every 1 hour at the wavelength of 600nm. Finally, the growth curve of the bacteria was drawn using the values obtained from the UV spectrophotometer

### Crystal violet staining

*S. xylosus* strains were grown in TSB for 24 h and diluted to a ratio of 1:100 in a fresh TSB which was pipetted into a sterile 96-well tissue culture plates. Then, 100μL aliquot of the cultures were added to each well of the 96-well microplate having equal volume of rutin already diluted to final concentrations of 0.2mg/mL, 0.4mg/mL, and 0.8mg/ml, respectively. A negative control (with TSB alone) and a positive control (with bacteria alone) were also included in the experimental set up. The cultures were incubated at 37°C for 6 h, 12 h and 24 h respectively, thereafter, the medium was removed by a micro pipette and the wells were washed three times with sterile phosphate buffer (PBS). The remaining attached bacteria were fixed with 200 μL of 99% methanol (Guoyao Ltd., China) per well, and the plates were emptied after 30 min. and left to dry. Afterwards, the plates were stained for 30 min. with 200 μL of 2% crystal violet (Guoyao Ltd., China) per well. The excess stain was rinsed out by placing the plate under running tap water. After the plates were air dried, the adherent cells were re-solubilized with 200μL of 33% (v/v) glacial acetic acid (Guoyao Ltd., China) per well. The amount of released stain was quantified by measuring the absorbance at 600 nm with a Microplate reader (DG5033A, Huadong Ltd., Nanjing, Jiangsu, China).

### Scanning Electron Microscope (SEM)

For SEM, *S. xylosus* strains were grown in TSB for 24 h at 37°C on the surfaces of sterile glass slides, which were deposited in advance in a 6 orifice plate having equal volume of rutin already diluted to a final concentration of 0.8mg/ml, at the same time, control groups (without rutin) were set up. SEM micrographs were taken in the electron microscopy core facility laboratory at the School of Life Sciences, Northeast Agricultural University.

### Molecular Docking of rutin and IGPD

Homology modeling of the IGPD of the *S. xylosus* and model validation have been described in our previous study [35]. The 3D structures of **rutin** was downloaded from Pubchem database (https://pubchem.ncbi.nlm.nih.gov/). The Ligand model that was implemented from the DS interface was used to optimize the rutin to generate different conformers for docking analysis in the Discovery Studio 3.0 (DS 3.0)”. We defined the two manganese ions of IGPD as the potential active pocket. Hence, other parameters were set to default. The best binding mode obtained was analyzed and visualized by PyMOL v2.3 software.

### Molecular dynamics simulation

Molecular Dynamics (MD) is one of the most commonly used methods in Molecular simulation, which is based on the mechanism of molecular force field. This method involves the use of the dynamic motion of molecules to describe the dynamic process of life. In our study, MD simulation was carried out on the receptor–ligand complex of the IGPD and rutin, using DS 3.0. Initially, we added a solvent environment of water to the protein to make the dynamics simulation closer to the real situation, salt concentration adopts a default value which was 0.145, cation type was Na^+^ and anionic type was Cl^-^. In addition, periodic boundary conditions were set up. The structural modifications and real time behavior of the complex structure were computed using root mean square deviation (RMSD) and potential energy of the receptor–ligand complex during the course of 2ms simulation time.

### Interactions of rutin and IGPD by SPR technique

The binding affinity of the rutin to the IGPD receptor, was investigated with a BIAcore 3000 SPR instrument (GE Healthcare, Little Chalfont, UK) using a CM5 sensor chip (GE Healthcare). This technique measures changes in the refractive index near the sensor surface [65]. IGPD was fixed i
n a flow cell on a sensor chip, and another cell was used as a reference. Afterwards, the other groups of active surfaces were blocked with 1 M ethanolamine, pH 8.5. The running buffer used for immobilization was 10 mM Hepes, 150 mM NaCl, 3.4 mM EDTA, pH 7.4. Interaction of IGPD on the surface was assayed in the presence of rutin at different concentrations. The mixture was injected with rutin (100μl) at a flow rate of 2 ml/min. All IGPD experiments were conducted at 25°C. Interaction of IGPD and rutin was measured as Resonance Units (RU)/s. The SPR signal obtained in each individual reaction cycle was recorded as a sensorgram, which is a real-time pattern plotted as RU versus time (s). After each experiment, the surface of the chip was regenerated by injection of 100μl inulinase from *Aspergillus* species (Sigma, St. Louis, MO, USA), 2.5 units/ml and 100μl of 5 mmol NaOH at flow rate of 0.5 ml/min. Data are expressed as relative RU.

### Effect of rutin on IGPD activity

*S. xylosus* was were grown in TSB for 6, 12 and 24 h respectively with 0.8mg/ml rutin, no rutin was added to the control groups. After culturing, for 20 mins, 5 mL of the bacterial solution was centrifuged (12 000 r/min, 2 min), the supernatant was discarded, the pre-conFigd PBS buffer solution was added, the cells were suspended, the bacteria were broken by ultrasound, then centrifuged again (12 000 r/min, 2 min), and the supernatant solution was retained as the sample for enzyme activity analysis. The activity of the enzyme was determined using a previously described stopped-assay protocol [66] with minor modifications. The reaction mixture consisted of PBS buffer pH 7.4, and IGPD. The reaction was carried out at 37°C using IGP (Santa Cruz Biotechnology, USA). The reaction was stopped by adding sodium hydroxide at the point with an interval of 30 s. The mixture was then incubated at 37°C for 20 min to convert the product imidazoleacetol-phosphate (IAP) into an enolized form, the absorbance was read at 280 nm using a Shimadzu UV spectrophotometer against a blank. The extinction coefficient of IAP formed under these conditions is as reported previously [66].

### Predicting the potential binding sites of rutin to IGPD by molecular docking

We analyzed the various binding states of rutin and IGPD. In the first four optimal binding modes, rutin and Ile27 and His36 can generate hydrogen bonds. Then we analyzed the bond lengths of these hydrogen bonds, which are also within a reasonable range. So, we suspect that these two amino acids may be the key amino acids that mediate the binding of rutin to IGPD. Subsequently the target amino acid was replaced by alanine by amino acid allele substitution, and the 3D structure of three mutant proteins (Ile27 mutant protein, His36 mutant protein and Ile27 and His36 double mutant protein) were obtained. The binding of rutin to three mutant proteins was analyzed by molecular docking. The experimental method was consistent with that of rutin and IGPD.

### Site direct mutagenesis, expression and purification of the IGPD mutant

The gene (*hisB*) encoding for IGPD from *S. xylosus* was designed and cloned into pET30a vector (Novagen), as pET30a-*hisB*. The pET30a-*hisB* mutant was made by introducing a point mutation (lle27, His36; double lle27 and His36) via Quick Change site-directed mutagenesis (TAKARA) and subsequently verified by sequencing (BGI). The vectors were subsequently cloned into a pET30a vector for protein expression and verified by PCR. Plasmids encoding for IGPD were transformed into *Escherichia coli* BL21 (DE3) cells (Novagen) and protein expression was induced for 5 h with 1 mM isopropyl-b-D-thiogalactopyranoside at 37°C in Luria broth. Cells were harvested and lysed by sonication. For IGPD, insoluble materials were removed by centrifugation and the cell-free extract was purified in a process involving affinity chromatography and molecular sieve chromatography. All proteins were analyzed after each stage by SDS-PAGE.

### Effect of rutin on the activity of IGPD mutant

After adjusting the protein concentration, rutin with the final concentration of 0.8mg/ml was added and control group was set up in a similar manner but without rutin. The analytical method was consistent with that of rutin and IGPD. The reaction was conducted at 37°C using IGP (Santa Cruz Biotechnology, USA). The reaction was stopped by adding sodium hydroxide at the point with an interval of 30 s. The reaction mixture was then incubated at 37°C for 20 min to convert the product imidazoleacetol-phosphate (IAP) into an enolized form, the absorbance was read at 280 nm using a Shimadzu UV spectrophotometer against a blank. The extinction coefficient of IAP formed under these conditions is as reported previously [66].

### Real Time RT-PCR

We investigated the effect of the 0.8 mg/ml of rutin on the gene expression of *hisB* in *S. xylosus*. So, it was grown with the 0.8 mg/ml of rutin at 37°C for 6, 12 and 24 h respectively.

Briefly, bacteria strains were collected by centrifugation (10,000 g for 5 min) and treated with an RNASE REMOVER (Huayueyang Ltd, Beijing, China). Total RNA was extracted with a bacterial RNA isolating kit (Omega, Beijing, China) and then, reverse transcription of total RNA into cDNA was carried out using a reverse transcription Kit. The reverse transcription conditions were set at 37°C for 15min and 85°C for 5s. In addition, Real-time PCR was used to measure the expression level of *hisB* mRNA. Relative copy numbers and expression ratios of selected genes were normalized to the expression of *16S-rRNA* gene (housekeeping gene). Besides, the specific primers used for the fluorescent quantitative PCR were designed by Bao Biological Corporation and are listed in Table 1. Finally, triplicate reactions were prepared with 10μL of PCR mixture consisting of 1μL cDNA, 0.6μL specific primers, 3.4μL dH_2_O, and 5μL roche dye, respectively in a 25μL IQ SYBR Green Supermix and the reaction was carried out on an ABI 7500 Real Time PCR System. The reaction conditions were as follows: 1 min at 95°C, 30s at 95°C, 30s at 60°C and 40 cycles [67].

**TABLE 1.**
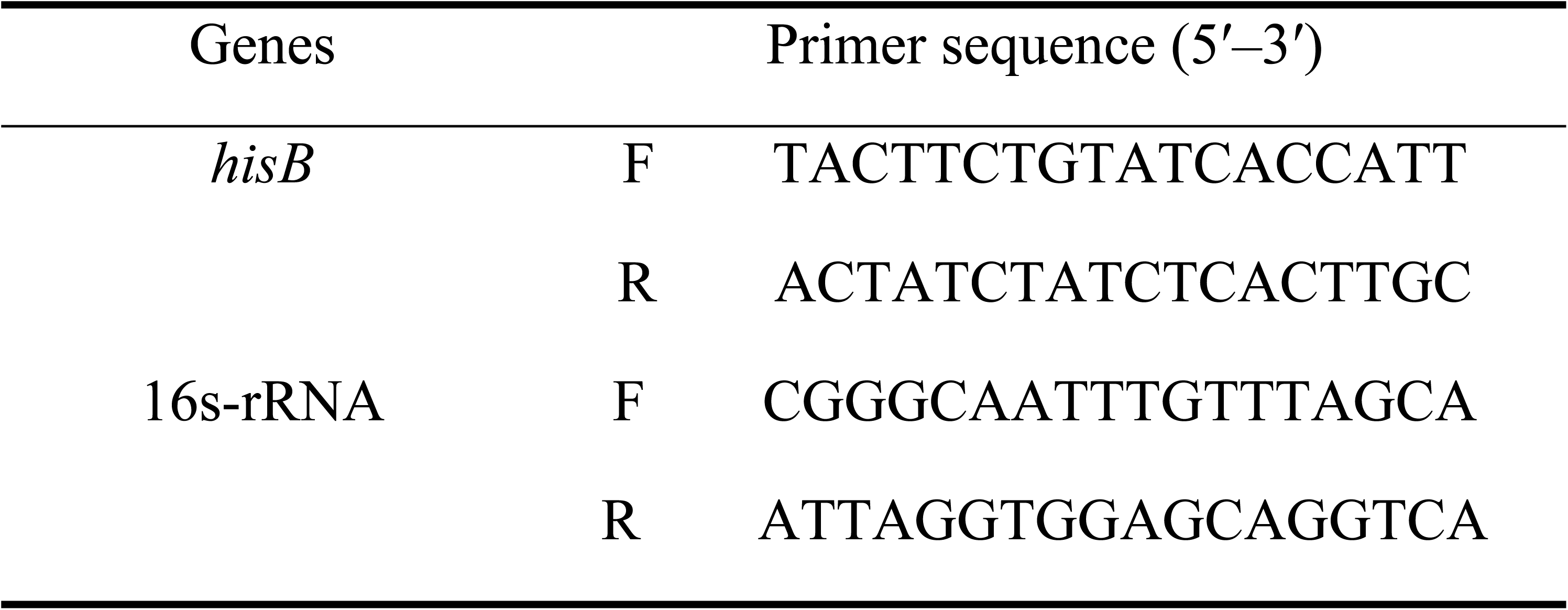
Primers used for the quantitative RT-PCR analysis.

### Western Blot

We investigated the effect of the 0.8 mg/ml of rutin on the expression of IGPD in *S. xylosus*. So, it was grown with the 0.8 mg/ml of rutin at 37°C for 6, 12 and 24 h respectively. Appropriate amount of PBS buffer solution was added to the harvested bacterial strains, and the solution was subjected to ultrasonic screening. This was followed by the centrifugation of the supernatants to obtain protein samples.

The protein samples were separated by 8% and 12% SDS PAGE and were transferred to PVDF membranes (Merck Millipore, USA, Cat# ISEQ. 00010, LOT#R6PA4145H). The membranes w ere blocked with 5% skim milk for 3 h at 37 °C and were incubated for 14 h at 4 °C, with the sel ected diluted primary antibodies. After washing three times for 15 min each with PBST, the mem branes were incubated for 2 h at 37 °C with peroxidase conjugated secondary antibodies against rabbit IgG (Santa Cruz Biotechnology, Argentina, Cat# sc2357, RRID: AB_628497). After washi ng thrice for 15 min each, the bound antibodies were visualized by chemiluminescence using the ECL plus reagent (GE Healthcare, Buckinghamshire, UK). The GAPDH content was analyzed as the loading control using a rabbit polyclonal antibody.

### Determination of histidine content

*S. xylosus* and the *ΔhisB S. xylosus* were grown with 0.8 mg/ml rutin at 37°C for 6, 12, and 24h, respectively. The pellets were then suspended in sterile double distilled water and the bacterial culture was sonicated to release histidine. The mixture was filtered using a 0.45 μm filter, and the histidine content in the bacteria was determined by high performance liquid chromatography (HPLC) on a Waters Alliance HPLC system (Waters e2695, United States). The stan dard solution of 15 mg histidine (99% pure histidine was purchased from Beijing Solarbio Technology Co., Ltd.) was prepared by dissolving it in 250 mL (0.1 mol/L) hydrochloric acid. The determination was performed on a 5 μm Diamosil C18 column (4.6 mm × 15 0 mm, Japan). The chromatographic separation was carried out on a Diamosil C18 column (4.6 mm × 250 mm, 5 μm) with a gradient solvent A (10 mmol/L diammonium hydrogen phosphate buffer (containing 10 mmol/L sodium 1octanesulfonate and obtaining pH 2.0 by phosphoric acid) and solvent B (acetonitrile) as mobile phase at a flow rate of 1 ml/min. The gradient con ditions were 0 to 5 min, 95% solvent A; 5 to 6 min, 95% to 86% solvent A; 6 to 15 min, 86% sol vent A; 15 to 16 min, 86% to 87% solvent A; 16 to 25 min, 87% solvent A. The detection wavelength was set at 205 nm, and the injection volume was 100 μL. The column temperature was set a t 8°C. Quantification of histidine in the sonicated treated bacteria was done in HPLC at 205 nm against concentration using the external standard method.

### Statistical Analysis

Assays were done three times and the means ± standard deviations were computed. Data were analyzed using the Student-test. (\**p* <0.05).

